# Thrombospondins 1 and 2 affect lysyl oxidase protein and collagen matrix maturation in cortical bone of growing male and female mice via non-redundant pathways

**DOI:** 10.1101/287763

**Authors:** Dylan Shearer, Madison O Mervis, Eugene Manley, Anita B Reddy, Andrea I Alford

**Author notes:** To whom correspondence should be addressed: Andrea I. Alford, Department of Orthopaedic Surgery, University of Michigan School of Medicine, A. Alfred Taubman Biomedical Sciences Research Building, Room 2009, Ann Arbor, MI, 48109.

## Abstract

Thrombospondin-2-deficiency is associated with impaired matrix maturation in osteoblasts and cortical bone of growing mice. Here we addressed the possibility that lysyl oxidase (LOX) contributes to this phenotype. After overnight serum starvation, pro-LOX levels were elevated compared to wild-type in marrow-derived osteoblasts from male and female TSP2−/− mice. The liberated LOX pro-peptide (LOPP) was faintly visible in serum-starved cultures. When serum was maintained, pro-LOX content was not affected by TSP2 status, but relative LOPP levels were elevated in cultures from female TSP2−/− mice. Two isoforms of pro-LOX at 75 kDa and 50 kDa were detected in detergent soluble protein extracts of diaphyseal tissue from growing mice. In female mice, TSP2 status did not affect detergent soluble pro-LOX content or the relative contribution of each band to the total signal. Instead, levels of the 50 kDa band were reduced in female TSP1−/− samples. In male diaphyseal tissue, total pro-LOX content and the contribution each isoform made to the total signal was not affected by TSP1 or TSP2 status. We did not detect 32 kDa mature LOX in detergent soluble preparations of cells or whole bone tissue. Detergent insoluble hydroxyproline content was reduced in diaphyseal tissue obtained from female TSP1−/− and TSP2−/− mice. In male diaphyseal cortical samples, TSP2 but not TSP1 deficiency was associated with reduced insoluble hydroxyproline content. Our data suggest that the trimeric thrombospondins contribute to bone matrix quality via non-redundant mechanisms that are dependent on the unique tissue milieu of the male and female skeleton.

## Introduction

Thrombospondins (TSPs) are a family of secreted matricellular glycoproteins that modulate cell-cell and cell-matrix interactions. They bind structural extracellular matrix (ECM) proteins, as well as cytokines, growth factors and proteinases (1,2). The trimeric matricellular proteins, TSP1 and 2 have high sequence and structural homology, but they are relatively divergent in their amino terminal regions (3,4). Matricellular proteins are highly expressed during development and then re-expressed upon injury (5,6). One exception is bone, where TSP1 and TSP2 are both components of the ECM in growing and adult mice (7,8).

TSP1 and TSP2 have each been implicated in the regulation of collagen fibrillogenesis in soft connective tissues (9), and collagen is a crucial mechanical and biological component of the skeleton. When they are cultured under osteogenic conditions, primary marrow-derived mesenchymal stem cells (MSC) harvested from TSP2−/− mice assemble a mature, insoluble ECM with reduced total collagen content (10). In femoral diaphysis of growing TSP2−/− mice, collagen fibril morphology is altered and detergent soluble type I collagen content is elevated despite WT equivalent total hydroxyproline content(11). These data suggest that TSP2 facilitates collagen matrix maturation and point to the possibility of impaired collagen cross-linking in the TSP2-deficient osteoblast ECM.

Collagen crosslinking is mediated by lysyl oxidase (LOX). LOX is secreted in a pro-form (pro-LOX), which consists of an 18 kDa N-terminal pro-peptide (LOPP) and a 32 kDa mature domain (12). LOPP is glycosylated (13) in a tissue-specific manner, leading to additional molecular weight isoforms on immunoblots (12). Bone morphogenetic protein (BMP)-1 and related proteinases remove LOPP and thereby activate LOX (14). LOX catalyzes the oxidative deamination of the ε-amino group of lysine and hydroxylysine residues within the N- and C-terminal telopeptides of fibrillar collagens, leading to the formation of reactive aldehydes that covalently bond to lysine and hydroxylysine residues in neighboring collagen monomers to cause the formation of inter- and intra-strand crosslinks. These divalent crosslinks subsequently mature into trivalent crosslinks (15,16). LOX deficiencies are associated with decreased collagen fibril diameter, altered collagen fiber morphology and lathyrism (17–19). LOX−/− mice die perinatally and osteoblasts from these neonatal mice display impaired proliferation and differentiation in addition to their altered ECM (20).

The purpose of the experiments presented here was to address the possibility that LOX contributes to the osteoblast and bone phenotypes of growing TSP2−/− mice. We also sought to determine whether the TSP2 homolog, TSP1, contributes to the regulation of collagen matrix maturation in bone tissues of the growing murine skeleton. Our data suggest that TSP1 and TSP2 both impact collagen matrix maturation in the female skeleton, but that LOX likely only contributes to this phenotype in the TSP1−/− case. In the male skeleton, TSP2 (and not TSP1) appears to facilitate collagen matrix maturation via a LOX independent pathway.

## Materials and Methods

### Primary Antibodies utilized in Western blot

Pro-LOX and the liberated LOX pro-peptide (LOPP) were detected with a polyclonal antibody raised against residues 78-115 (Thermo Fisher catalog # PAI-16989). This peptide sequence is contained within the pro-peptide (21), so the antibody recognizes pro-LOX (50 kDa) and LOPP (18-32 kDa). Pro-LOX and mature LOX were detected using a polyclonal antibody raised against amino acids 200-300 of human LOX (Thermo Fisher catalog # PA1-16953). This peptide sequence is contained within the mature LOX domain, so the antibody recognizes pro-LOX (50-75 kDa) and mature LOX (32 kDa). The ranges of pro-LOX and LOPP apparent molecular weights is due to differential glycosylation of the pro-domain. Antibodies against α-actinin (Santa Cruz Biotechnologies) and β-actin (Novus Biologicals) were used as loading controls for cell matrix layers and bone protein extracts. Anti-cellular fibronectin was obtained from Sigma (F6140).

### Animals

All animal procedures were performed at the University of Michigan under Institutional Animal Care and Use Committee (IACUC) approval and comply with NIH guidelines outlined in the Guide for the Care and Use of Laboratory Animals. We maintain colonies of TSP1−/− (22) (Jackson Labs) and TSP2−/− (23) mice on the C57/B6 background. Animals were housed under specific pathogen free conditions and had free access to standard chow, water, and cage activity. 6-week old male and female mice were sacrificed by CO_2_ inhalation followed by bilateral pneumothorax. Animal numbers are noted in figure legends.

### Procurement of Skeletal Tissue

Tibiae and femora were dissected, cleaned of soft tissue with a Kim Wipe, flash frozen in liquid nitrogen and stored at −80°C. On the day of protein extraction, bones were cut over a liquid nitrogen-cooled chamber using a #10-scalpel blade. Proximal ends of femora were cut at the greater trochanter, and distal ends were separated from the mid-diaphysis. Tibiae were cut at the distal tibial-fibular junction and the proximal metaphysis was removed. Bone segments with marrow intact were immediately homogenized as outline below.

### Bone Protein Extraction

Detergent-soluble proteins were extracted as we have described (7,11). Briefly, bone tissues were crushed and homogenized on ice with a polytron in lysis buffer containing 50 mM Tris, 150 mM NaCl, 2mM EGTA, 10% NP40, 10 mM Na pyrophosphate decahydrate plus a protease inhibitor cocktail of 1mM PMSF, 5 mM sodium fluoride, 1 mM sodium orthovanadate and the Complete Protease Inhibitor Cocktail (Roche). Extracts were incubated 4 hours at 4°C with rocking. Samples were then centrifuged at 12,000 rpm for 20 minutes at 4°C. The supernatant was collected for western blot and the detergent-insoluble pellet was subjected to hydroxyproline analysis. Both fractions were stored at −80°C until the time of analysis.

### Primary cell culture

Primary marrow-derived mesenchymal stromal cell (MSC) culture was carried out as described (7,10,24). Whole marrow was flushed from femora and tibiae of 6-week old mice. A 23-gauge needle was used to create a single cell suspension, which was re-suspended in MSC growth medium (α-MEM containing 10% fetal bovine serum (FBS; Fisher Scientific, Pittsburgh, PA), 100 IU/ml penicillin, 100 μg/ml streptomycin, 2 mM L-glutamine (Glutamax; Invitrogen, Carlsbad, CA) and 50 μg/ml ascorbate-2 phosphate (Sigma-Aldrich, St. Louis, MO)). Cells were plated at a ratio of 1 mouse per 10 cm dish. A partial (1/3) medium change including fresh ascorbate was conducted on day 4. On day 11, non-adherent cells were discarded and the adherent MSC were passaged onto 6-well plates at 200,000 cells per well in MSC growth medium containing ascorbate 2-phosphate. Medium was changed every 3-4 days and fresh ascorbate 2-phosphate was added each time. When indicated, cells were serum starved 24 hours before harvest in medium containing 0.1% FBS. Cell-matrix layers were washed with cold PBS and then scraped into lysis buffer containing a cocktail of protease inhibitors in preparation for SDS-PAGE and Western blot.

### Western Blot

Detergent soluble bone protein extracts were diluted in gel loading buffer (62.5 mM Tris-HCl, pH 6.8, 2% SDS, 10% glycerol, 0.01% bromophenol blue and 5% β-mercaptoethanol).

Samples were boiled for 2-3 minutes, and separated on 12% (anti-LOPP, anti-LOX and anti-α-actinin) or 7.5%. (anti-fibronectin) SDS-PAGE gels. Proteins were transferred to nitrocellulose membranes in transfer buffer containing 50 mM Tris, 192 mM glycine, 0.1% SDS and 20% methanol. Non-specific sites were blocked overnight at 4°C with Tris-buffered saline containing 5% non-fat dry milk or 1% BSA (anti-LOPP) and 0.05% Tween 20. Blots were incubated overnight at 4°C with primary antibody and then washed with TBS-tween. Blots were then incubated with horseradish peroxidase conjugated secondary antibodies for 1 hour at room temperature. Membranes were washed in TBS-T and proteins were detected using enhanced chemiluminescence (ECL). Blots were sequentially re-probed with anti-α-actinin, anti-β-actin or anti-fibronectin. Band intensities were determined using an XRS+ Chemidoc system and Image Lab software (Biorad).

### Hydroxyproline Analysis

The insoluble pellets obtained above after detergent extraction were subject to hydroxyproline analysis to determine insoluble collagen content. Pellets were thawed, centrifuged at 10,000 rpm for 2 minutes to remove surface water, and wet weights were recorded. Pellets were then dried for 30 minutes at 110°C, hydrolyzed for 8 hours in 6N HCl at 130°C and neutralized with 6N NaOH. Hydroxyproline content was determined using standard techniques (25). One hundred microliters of neutralized samples and hydroxyproline standards were added in triplicates to a 96-well plate. Fifty microliters of 0.05 M chloramine T were added to each well and incubated for 20 min at room temperature. Perchloric acid (3.15 M) was then added, and plates were incubated for 5 min at room temperature. Ehrlich’s Solution (20%) was added for 20 min, and plates were incubated at 60°C. Absorbance was read at 560 nm using a UV SpectraMax M5 Plate Reader (Molecular Devices, Sunnyvale, CA). Hydroxyproline concentrations in the samples were determined using the standard curve and normalized to pellet wet weight.

### Statistics

Statistical analysis was conducted using Graph Pad Prism v7.0 (La Jolla, CA). Continuous data are expressed as mean and standard deviation. Differences between groups were identified using students t-test or ANOVA and Tukey post-hoc tests, as appropriate. Discrete data were analyzed by Chi square. Significance was set at p<0.05.

## Results

After overnight serum starvation, pro-LOX/α-actinin levels in detergent soluble cell-matrix preps were elevated compared to WT in TSP2−/− marrow-derived MSC cultured under osteoblastic conditions for 10 and 14 days. Similar results were obtained in cells from male and female mice (Figure 1 top panels). At day 10 pro-LOX/α-actinin values were 3.9±1.3 and 11.7±7.3 in female WT and TSP2−/− MSC, respectively (p=0.07). For the male samples, pro-LOX/ᾶactinin values were 2.9±1.4 and 6.5±2.1 in WT and TSP2−/− MSC, respectively at day 10 (p<0.01). At day 14, pro-LOX/α-actinin values were 3.0±0.8 and 7.3±3.3 in female WT and TSP2−/− MSC, respectively (p<0.05). For the male samples, pro-LOX/α-actinin values were 1.0±0.3 and 1.8±0.4 in male WT and TSP2−/− MSC, respectively at day 14 (p<0.01). Qualitatively similar results were obtained in Western blots of conditioned medium. Representative blots of cell-matrix layers and conditioned medium are shown in figure 1 (bottom panel).

**Figure 1.**
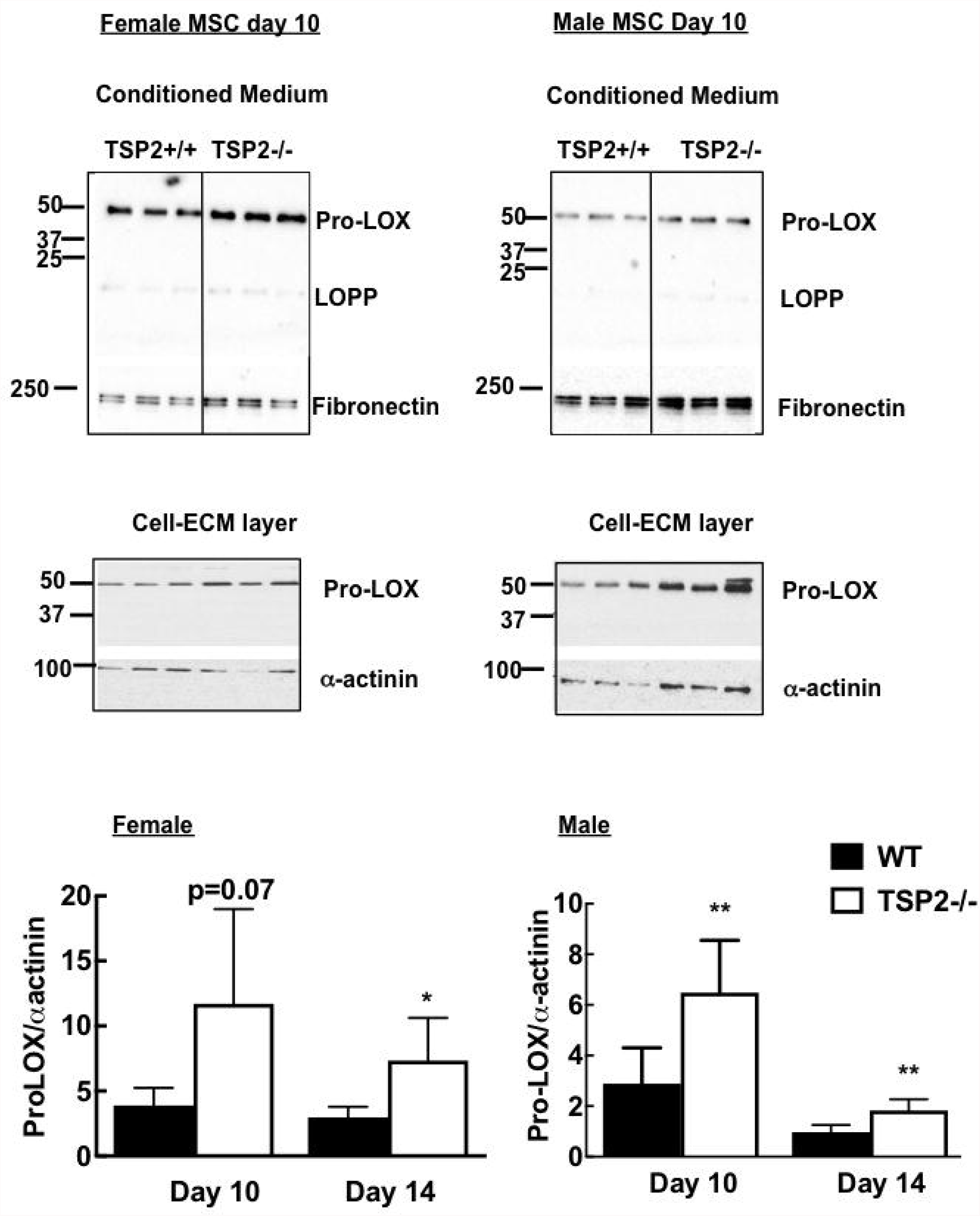
Elevated pro-LOX protein levels in primary cultures of marrow-derived osteoblast lineage cells from growing TSP2−/− mice. First passage MSC were cultured with ascorbate 2-phosphate to promote collagen matrix formation. Twenty-four hours before the indicated harvest times, cells were starved in medium containing 0.1% FBS. Pro-LOX content in medium (equal volume) and detergent soluble cell-matrix layers (equal protein) was determined by Western blot using an antibody against a peptide in the pro-domain. Data from female and male cultures are presented on the left and right side, respectively. Top panels: Representative blots of conditioned medium probed for pro-LOX and fibronectin. Middle panels: representative blots of cell matrix layers probed for pro-LOX and α-actinin. Bottom panels: Bars are mean and SD of proLOX/α-actinin for female (left panel; n=4 wells per genotype and time point) and male (right panel; n=6 wells per genotype and time point). Two primary cell harvests were conducted per sex and genotype. * p<0.05, ** p<0.01 vs. WT at the same time point.

Fibronectin content was not significantly affected by TSP2-status after 24 hours serum deprivation. At day 10, medium fibronectin/cell α-actinin values were 0.61±0.2 and 2.5±2.7 in WT and TSP2/-, respectively in female cells and 5.79±4.3 and 8.71±2.6 in WT and TSP2−/−, respectively in male cells. At day 14, fibronectin/α-actinin values were 0.86±0.6 and 1.87±0.6 in WT and TSP2−/−, respectively in female cells and 2.65±1.2 and 3.63±1.8 in WT and TSP2−/−, respectively in male cells.

The anti-LOX propeptide antibody used in figure 1 detects pro-LOX (50-75 kDa) and the liberated LOX-pro-peptide (LOPP; 18-32 kDa) (26). When the cells were serum starved overnight, LOPP was faintly visible only in conditioned medium of female cells. Supplemental figures S1 and S2 show complete representative blots with molecular weight markers for the serum starved cell culture experiments.

When we omitted the 24-hour serum starvation step, LOPP accumulated in the conditioned medium and we were able to detect it (Figure 2). As a surrogate marker of LOX activation, we analyzed relative LOPP content in conditioned medium by summing the band intensity values for pro-LOX and LOPP and then determining the percentage of this total that was due to LOPP. The proportion of total LOX signal due to LOPP was elevated in TSP2−/− cultures (Figure 2A). Thus, at day 10, 30.4±24 and 62.5±20.4 percent of the total LOX signal was due to LOPP in WT and TSP2−/− cultures, respectively (p<0.05). At day 14, 53.9±16.6 and 77.6±15.3 percent of the total LOX signal was due to LOPP in WT and TSP2−/− cultures, respectively (p<0.05). In contrast to results shown in figure 1, pro-LOX levels in the cell matrix layer were not affected by TSP2 status when the serum starvation step was omitted (Figure 2B). We also detected additional larger isoforms of pro-LOX that likely represent differentially glycosylated isoforms. LOPP was barely detectable in a few detergent soluble cell-matrix layers (Supplemental Figure 4). Similarly we were unable to detect mature LOX (32 kDa) in medium or detergent soluble cell-matrix layers using the anti-LOX antibody (data not shown). Medium fibronectin content per cell layer β-actinin was not affected by TSP2 status at day 10 (2.27±1.0 in WT and 4.36±3.2 in TSP2−/− samples) or at day 14 (4.38±3.2 in WT and 4.0±2.9 in TSP2−/− samples). Supplemental figures S3 and S4 show complete representative blots with molecular weight markers for the serum-replete cell culture experiments.

**Figure 2.**
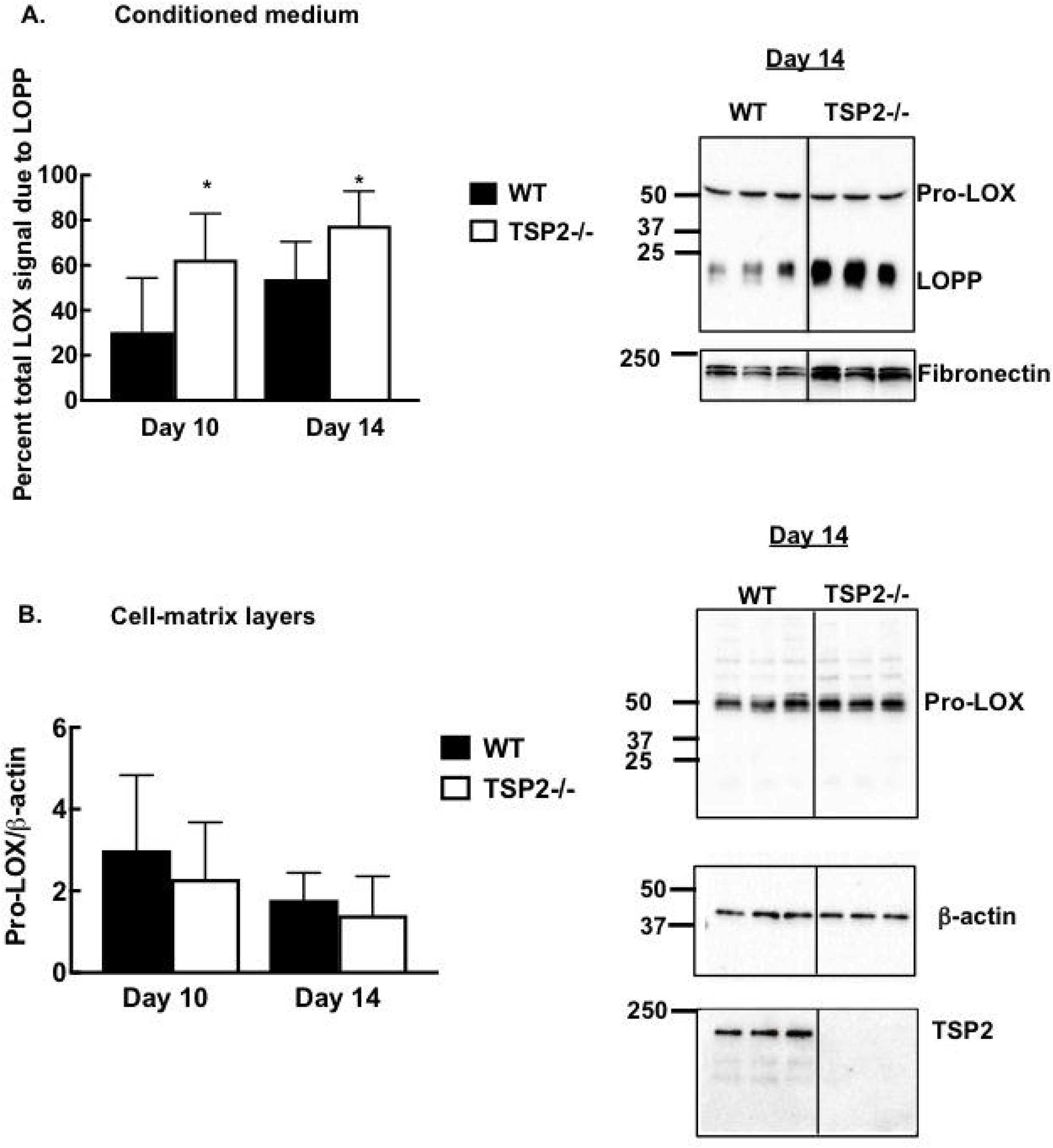
Elevated LOPP levels in primary cultures of marrow-derived osteoblast lineage cells from growing female TSP2−/− mice. First passage MSC were cultured with ascorbate 2-phosphate to promote collagen matrix maturation. Conditioned medium (A) and cell-matrix layers (B) were collected at the indicated times without prior serum starve. A (left panel): bars are mean and SD of LOPP band intensity expressed as a percentage of total LOX band intensity (pro-LOX + LOPP). A (right panel): Representative blots of cell-conditioned medium from day 14 samples are shown. B (left panel): Detergent-soluble cell-matrix layers corresponding to the medium collected in A were subjected to Western blot using anti-pro-LOX, anti-β-actin and anti-TSP2 antibodies. Bars are mean and SD of pro-LOX band intensity normalized to that of 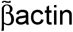. B (right panel): Representative blots of day 14 cell-matrix layers are shown. Data are from 3 independent cell harvests totaling 4-6 wells per time point. *p<0.05 vs. WT at same time point.

Next, detergent soluble protein extracts prepared from femoral diaphysis of growing (6-week old) female and male mice were subjected to Western blot with an anti-LOX antibody that recognizes pro-LOX (50-75 kDa) and mature LOX (32 kDa). We included both TSP1−/− and TSP2−/− samples in this experiment to address potential functional redundancy of these matricellular proteins. In female diaphyseal tissue, two immunoreactive species were detected at approximately 50 kDa and 75 kDa. These are likely differentially glycosylated isoforms of pro-LOX (12). In detergent soluble protein extracts of whole bone tissue, we did not detect a band near 32 kDa, where mature LOX with its pro-peptide removed is expected to run. Total detergent soluble pro-LOX content was not significantly affected by TSP1 or TSP2 status (figure 3A, middle panel). Values of total LOX/α-actinin were 4.52±3.7, 2.55±2.0 and 2.58±2.5 in WT, TSP1−/− and TSP2−/− samples, respectively.

**Figure 3.**
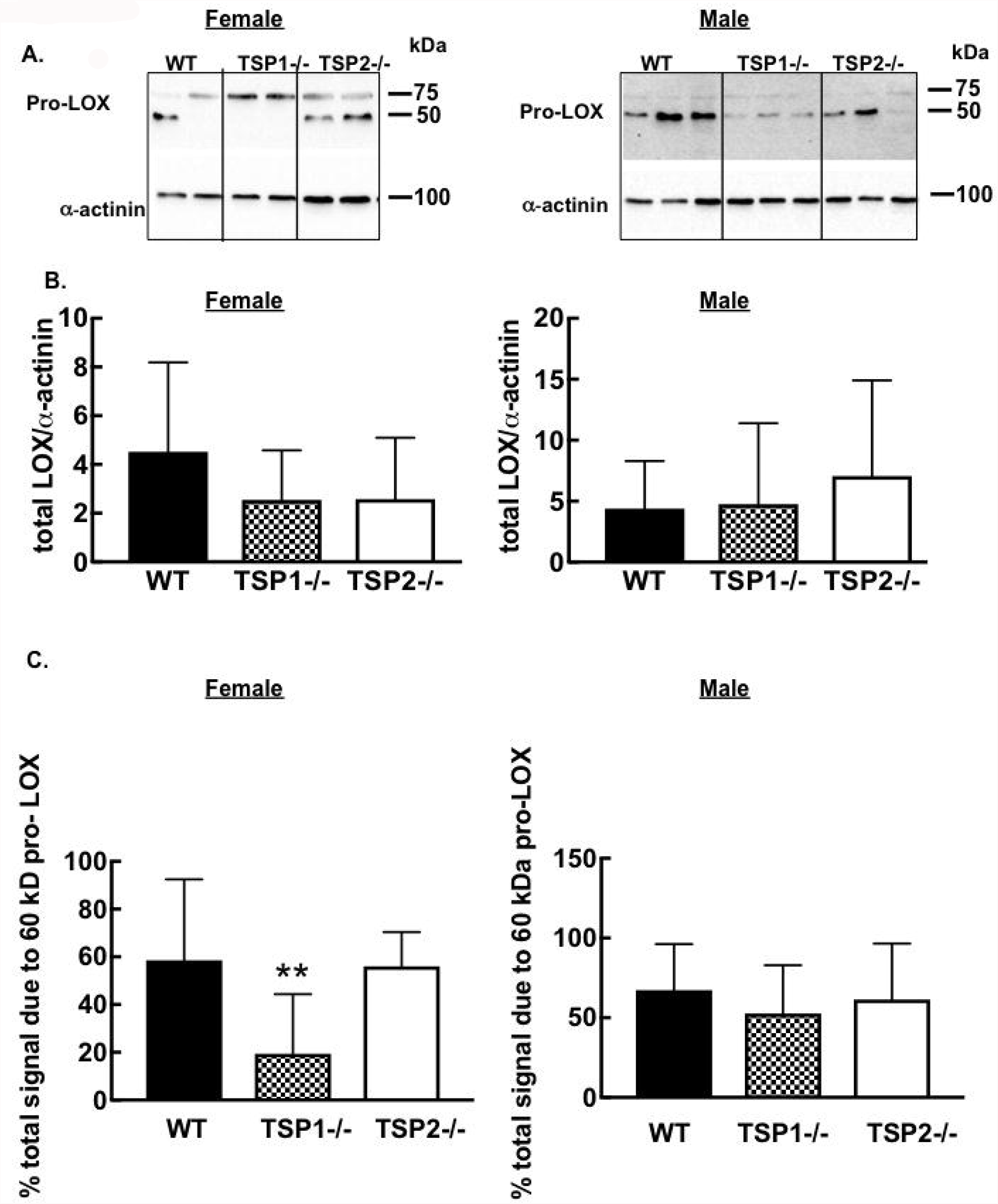
Relative levels of detergent soluble pro-LOX isoforms are affected by TSP1 status in femoral diaphysis of growing female mice. **A.** Detergent soluble protein extracts were prepared from femoral diaphysis of female (left) and male (right) mice and subjected to Western blot with anti-LOX and anti-α-actinin antibodies. **B.** Bars are mean and SD of total detergent soluble pro-LOX content (75 kDa + 60 kDa pro-LOX/α-actinin). **C.** Bars are mean and SD of 60 kDa pro-LOX band intensity expressed as a percentage of total LOX intensity. *p<0.05 by ANOVA. For female samples, N= 13 WT, 8 TSP1−/− and 14 TSP2−/− mice. For male samples, N=9 WT, 8 TSP1−/− and 6 TSP2−/− mice.

Instead, the relative contribution that each pro-LOX species (75 kDa and 50 kDa) made to the total pro-LOX signal varied according to TSP status (Figure 3A, bottom panel). 60 kDa pro-LOX accounted for approximately 60% of the total detergent soluble LOX signal in both WT and TSP2−/− groups, while it was significantly reduced in the TSP1−/− group. Thus, the lower 60 kDa molecular weight isoform of pro-LOX accounted for 58.51±34.0, 19.39±25.0, and 56.04±14.3 percent of the total LOX signal in WT, TSP1−/− and TSP2−/− samples, respectively (p<0.01 TSP1−/− vs. WT; p<0.05 TSP1−/− vs. TSP2−/−).

When the same analysis was conducted on diaphyseal tissue obtained from growing male mice, the same immunoreactive species were detected. TSP status again did not affect total detergent soluble pro-LOX content. Thus, total pro-LOX per αactinin values were 4.39±3.9, 4.75±6.6 and 7.05±7.9 in WT, TSP1−/− and TSP2−/− samples, respectively (Figure 3A, right panel). The percentage of the total detergent soluble LOX signal that was due to the 60kDa isoform of pro-LOX in male diaphyseal tissue was not affected by TSP status (Figure 3B, right panel). The 60 kDa pro-LOX band contributed to 67.32±28.8, 52.66±30.3 and 61.4±35.2 percent of the total LOX signal in WT, TSP1−/− and TSP2−/− samples, respectively.

In this study, visibility of the 60 kDa pro-LOX band was highly variable in the protein extracts prepared from female bones. Supplemental Figures S5-S8 show complete blots with molecular weight markers for all diaphyseal tissue analyzed in this paper. In select samples, the 60 kDa band was not visible. Thus, the 60 kDa band was visible in 9 of 12; 3 of 9 and 11 of 11 female WT, TSP1−/− and TSP2−/− samples, respectively (p<0.01 by chi square). For the male samples, the 60 kDa band was visible in 9 of 10; 7 of 9 and 6 of 7 WT, TSP1−/− and TSP2−/− respectively (not significant by chi square).

Since we were unable to detect 32 kDa mature LOX in our detergent soluble whole bone protein extracts, we used the anti-LOX pro-peptide antibody to address the possibility that LOX activation might be affected by TSP status. Figure 4A shows that LOPP was visible in all samples, and its contribution to the total lysyl oxidase signal was roughly equivalent in all groups (Figure 4).

**Figure 4.**
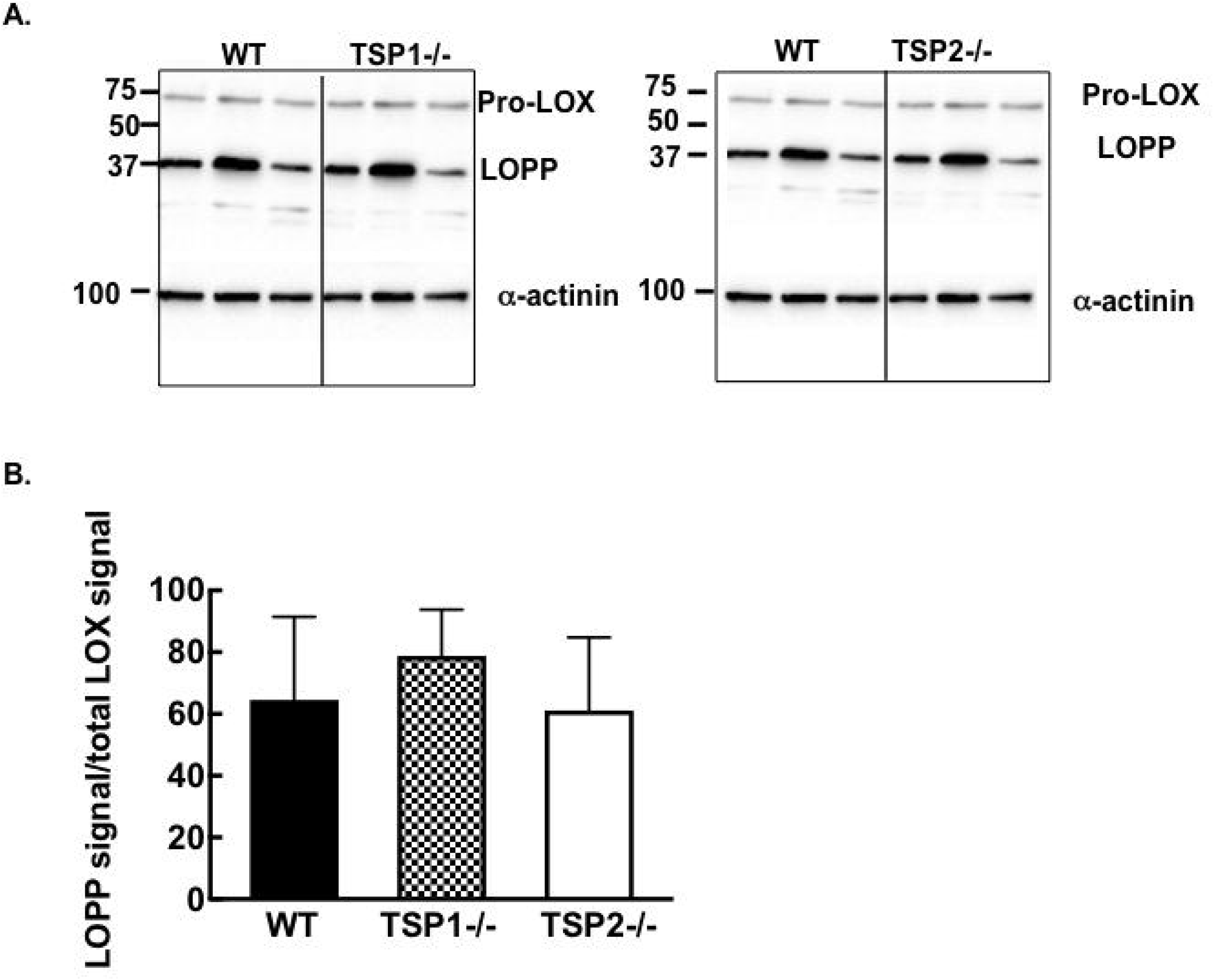
Removal of the LOX pro-peptide in femoral diaphysis of growing female mice is not affected by TSP1 or TSP2 status. **A.** Detergent soluble protein extracts were prepared from femoral diaphysis of growing female mice and subjected to Western blot with anti-pro-LOX and α-actinin antibodies. **B.** Bars are mean and SD of LOPP band intensity normalized to that of total LOX content (pro-LOX+LOPP) and expressed as a percentage. N=10 WT, 4 TSP1−/− and 9 TSP2−/− mice.

To determine the extent to which altered post-translational modification of detergent soluble pro-LOX might be associated with matrix maturation, we measured hydroxyproline content in detergent insoluble pellets obtained from diaphyseal cortical bone. Femoral diaphyseal tissue from both TSP1−/− and TSP2−/− female mice displayed significant reductions in detergent-insoluble hydroxyproline content (Figure 5A). Values for μg hydroxyproline per mg insoluble pellet wet weight were 1.70±0.5, 1.00±0.3 and 0.96±0.1 in WT, TSP1−/− and TSP2−/− samples, respectively (p<0.05).

**Figure 5.**
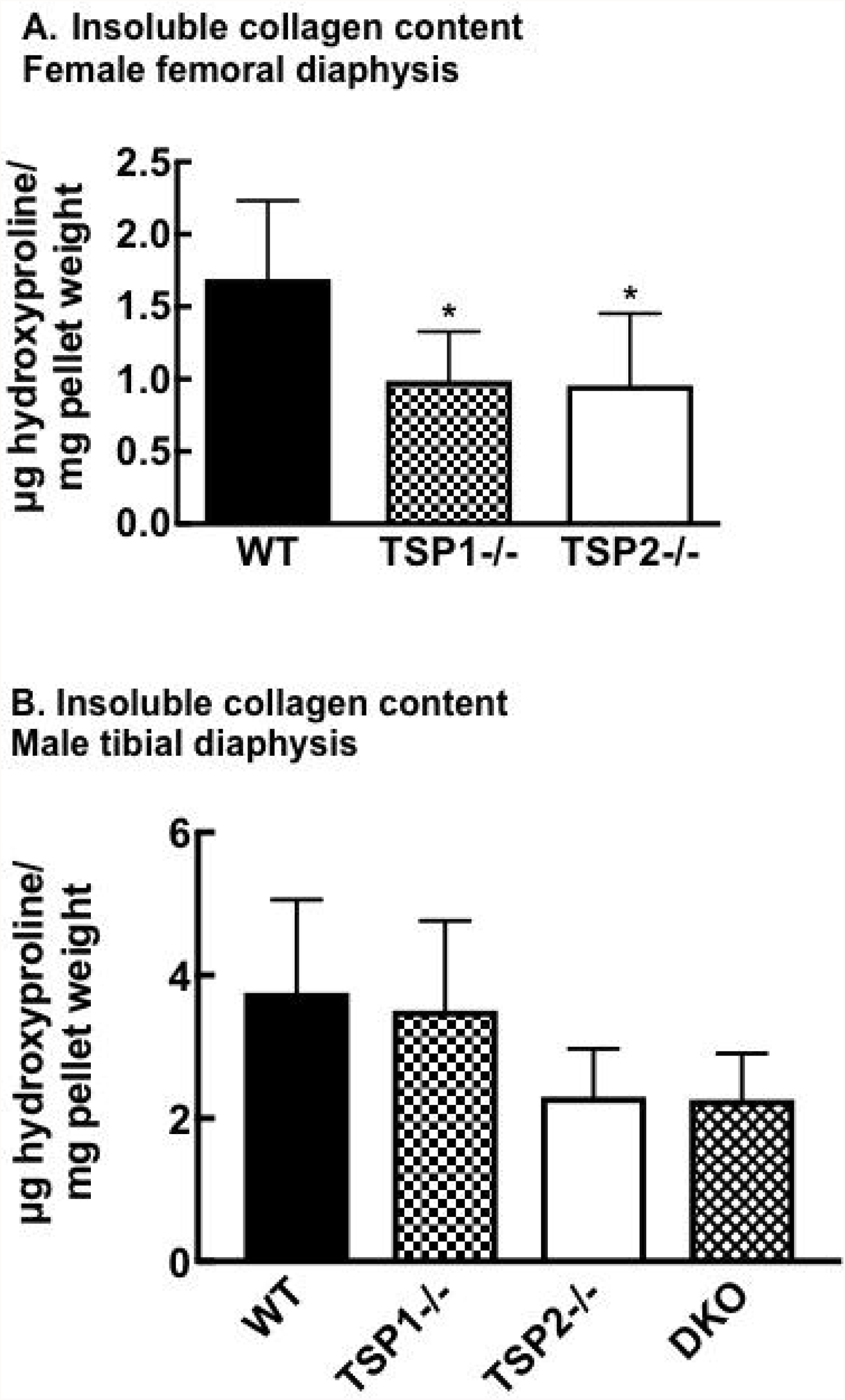
TSP1 and TSP2 both contribute to collagen matrix maturation in the growing female skeleton, but TSP2 plays a dominant role in the male skeleton. Bars are mean and SD of insoluble hydroxyproline content of diaphyseal cortical bone obtained from female (A) and male (B) mice. * p<0.05 compared to WT. For female mice N=7 WT, 7 TSP1−/− and 6 TSP2−/− insoluble pellets. For male mice, N=6 WT, 6 TSP1−/−, 6 TSP2−/− and 6 TSP1-TSP2 DKO insoluble pellets.

In male cortical bone samples, TSP2-deficiency was associated with modest decreases in total insoluble collagen content, but TSP1 status did not affect this outcome measure (Figure 5B). The effect of TSP2-deficiency was equivalent in TSP2−/− and TSP1-2 double knockout samples. Thus, values for μg hydroxyproline per mg insoluble pellet wet weight were 3.76±1.3, 3.51±1.3, 2.31± 0.7 and 2.26±0.6 in WT, TSP1−/−, TSP2−/− and TSP1-TSP2 double knockout tibial diaphysis respectively (ANOVA p<0.05; p=0.09 for TSP2−/− vs. WT; p=0.08 for DKO vs. WT). Femoral diaphysis from TSP2−/− male mice also displayed decreased insoluble hydroxyproline content, but the difference did not reach significance by ANOVA (1.08± 0.8, 1.12± 0.8 and 0.53±0.2 in WT, TSP1−/− and TSP2−/− samples, respectively).

## Discussion

We used Western blot analysis to examine pro-LOX in osteoblasts and whole bone tissue derived from growing male and female WT, TSP1−/− and TSP2−/− mice. We detected immunoreactive isoforms with molecular weights corresponding to differentially glycosylated isoforms of pro-LOX and LOPP. We did not detect the expected 32 kDa mature LOX band and we hypothesize that active LOX is present in the mature insoluble collagenous ECM. Since LOPP was detected in all of our detergent soluble extracts of whole bone, we think LOX pro-peptide removal is not affected by TSP status. Others have detected a 32 kDa isoform corresponding to mature LOX in cell culture (13) and the nature of this discrepancy is not understood currently. Nevertheless, glycosylation of the pro-domain is known to impact LOX enzyme activity and so we cannot rule out the possible contributions of TSP1 to LOX function (27).

Data presented here suggest that TSP2-dependent regulation of LOX contributes, at least in part to the phenotype of TSP2−/− osteoblasts and bone tissue (10,11). Thus, LOPP accounted for a higher percentage of total soluble LOX content in TSP2−/− cultures. More robust removal of the LOX-pro-peptide suggests increased LOX activation under conditions of TSP-2 deficiency. However, this possibility seems unlikely given that TSP2−/− osteoblasts and bone ECM display reductions in insoluble cross-linked collagen content (Figure 5), as well as deficits in collagen fibrillogenesis and matrix maturation (10,11). Alternatively, TSP2 might affect the fate or function of the liberated LOPP (26,28) or facilitate functional interactions between mature active LOX and collagen. We were not able to detect mature LOX with its pro-peptide removed in conditioned medium or detergent soluble cell layers. One possibility is that these elements are incorporated into the mature, cross-linked detergent insoluble ECM formed during in vitro osteoblast matrix maturation.

Our results suggest that TSP2-deficiency enhanced secretion or stability of pro-LOX during a 24-hour serum-starvation period. This effect of TSP2 on pro-LOX was lost when cells were cultured in the presence of 10% FBS and medium was changed every three days. This observation suggests that factors in serum might help normalize pro-LOX content by limiting TSP2 interactions with pro-LOX. Alternatively, serum deprivation and add-back regulates LOX gene expression in rat aortic smooth muscle cells (12). We think TSP2-dependent regulation of LOX gene expression under standard osteoblast differentiation conditions in vitro or in cortical bone tissue in situ is less likely, because pro-LOX (50 kDa) levels were not affected in TSP2−/− cells cultured with 10% FBS or in TSP2−/− diaphyseal tissue.

TSP1 and TSP2 are secreted modular glycoproteins that form 450 kDa homotrimers and they share substantial homology (3). TSP1 and TSP2 have each been implicated in the regulation of collagen fibrillogenesis (9-11,29), so we examined tissue from mice deficient in each TSP to address potential functional redundancy with regard to collagen matrix maturation. Our data suggest that TSP1 and TSP2 facilitate matrix maturation in femoral cortical bone of female mice via non-redundant mechanisms. In female diaphyseal cortical tissue, TSP1 and TSP2-deficiency both impacted insoluble hydroxyproline content, a marker of collagen crosslinking and thus matrix maturation. The mechanisms leading to impaired matrix maturation in the female TSP1−/− and TSP2−/− diaphysis appear to be different. TSP1-deficiency did not impact detergent soluble type I collagen content as determined by Western blot (data not shown). This is in contrast to the TSP2−/− diaphyseal bone tissues, in which detergent soluble type I collagen content is elevated despite WT equivalent total hydroxyproline content (11). Thus the collagen types representing total insoluble hydroxyproline content might be differentially modulated by TSP1 and TSP2-deficiencies.

Total detergent soluble pro-LOX content was not affected by TSP status in diaphyseal cortical tissues harvested from male or female mice. Instead, the relative contribution of the smaller (50 kDa) isoform to the total pro-LOX signal was significantly reduced in female (and not male) TSP1−/− samples. This result was associated with the observation that the 50 kDa isoform was not visible in a significant proportion of the female TSP1−/− diaphyseal tissue extracts. These data suggest that TSP1-deficiency affects post-translational modification of pro-LOX in the female skeleton.

Taken together, our data suggest that the TSP1 and TSP2 regulate collagen matrix maturation via mechanisms that are distinct in the skeletons of growing male and female mice, and thereby contribute to bone matrix quality.

