## Supplementary figures and images for "Thrombospondins 1 and 2 affect lysyl oxidase protein and collagen matrix maturation in cortical bone of growing male and female mice via non-redundant pathways"

### Supplementary Materials

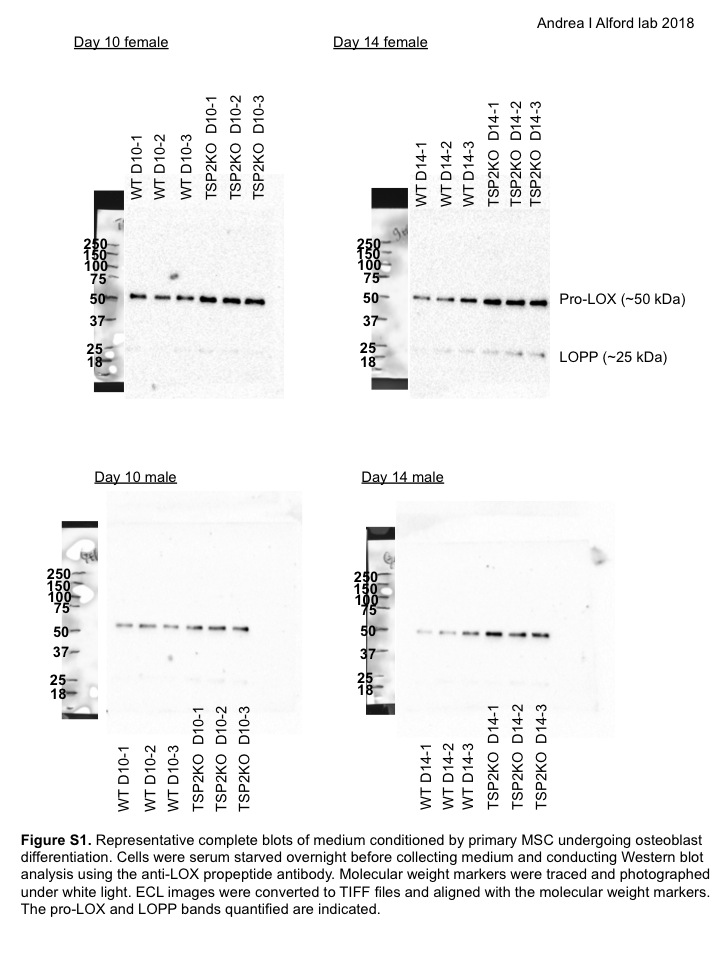

### Supplementary Materials

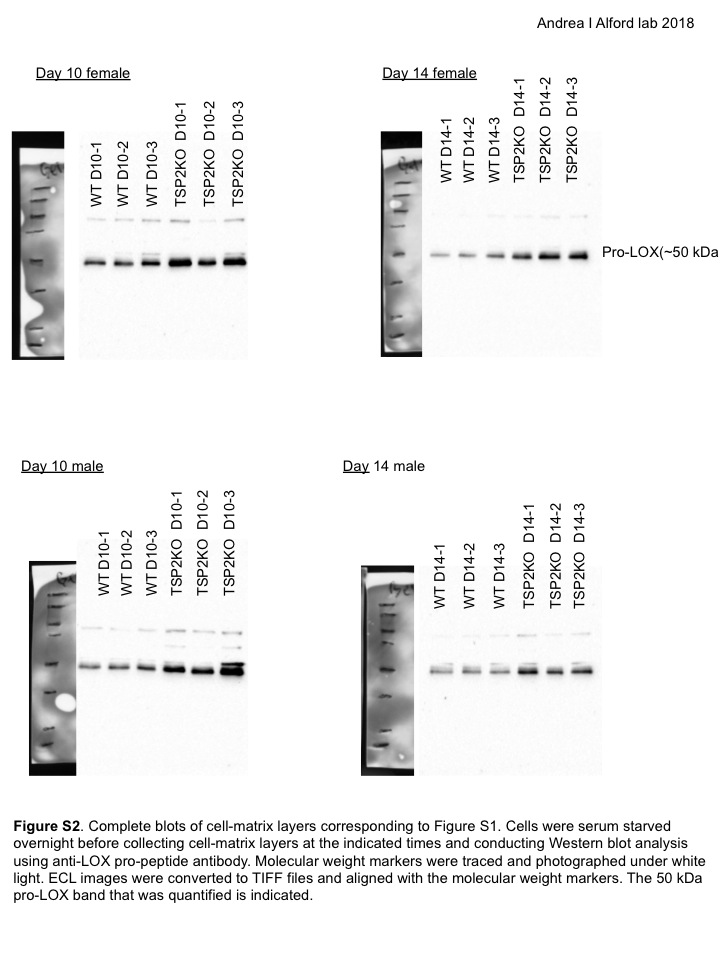

### Supplementary Materials

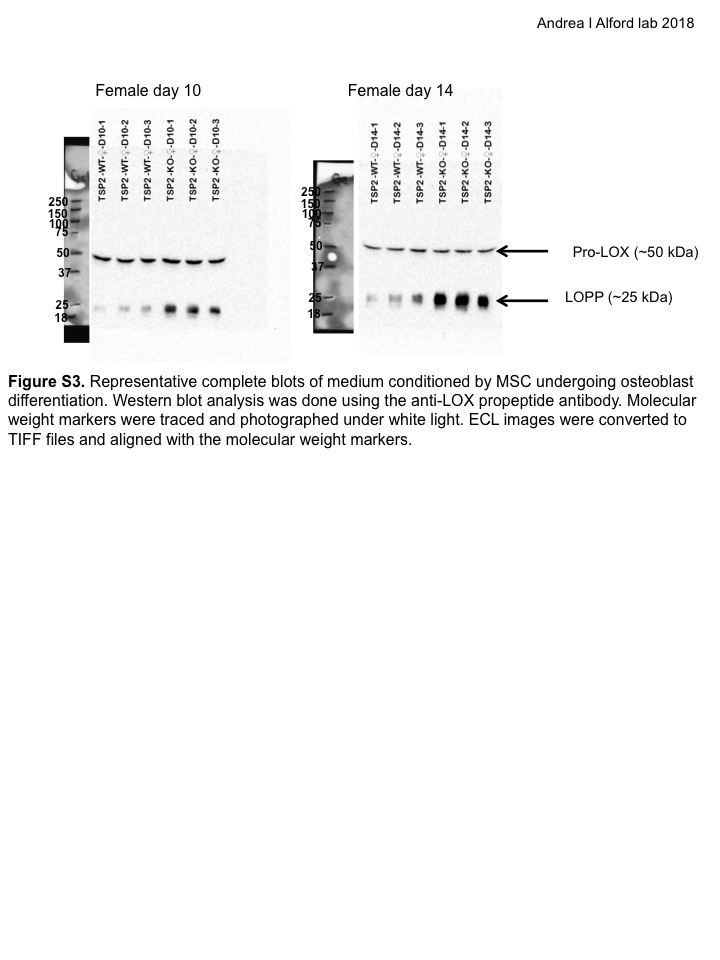

### Supplementary Materials

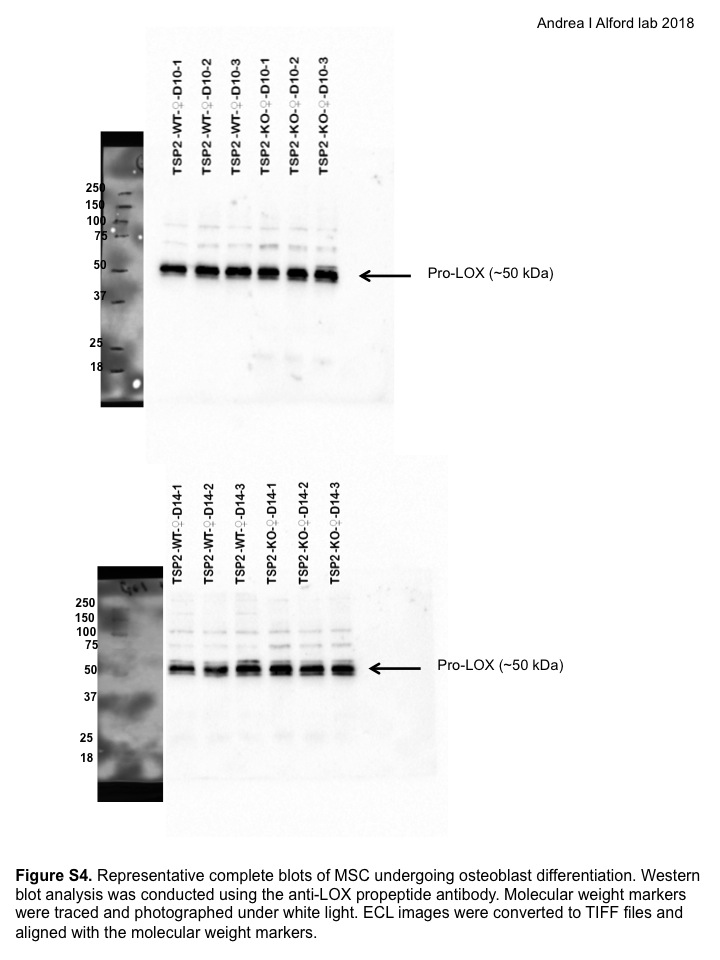

### Supplementary Materials

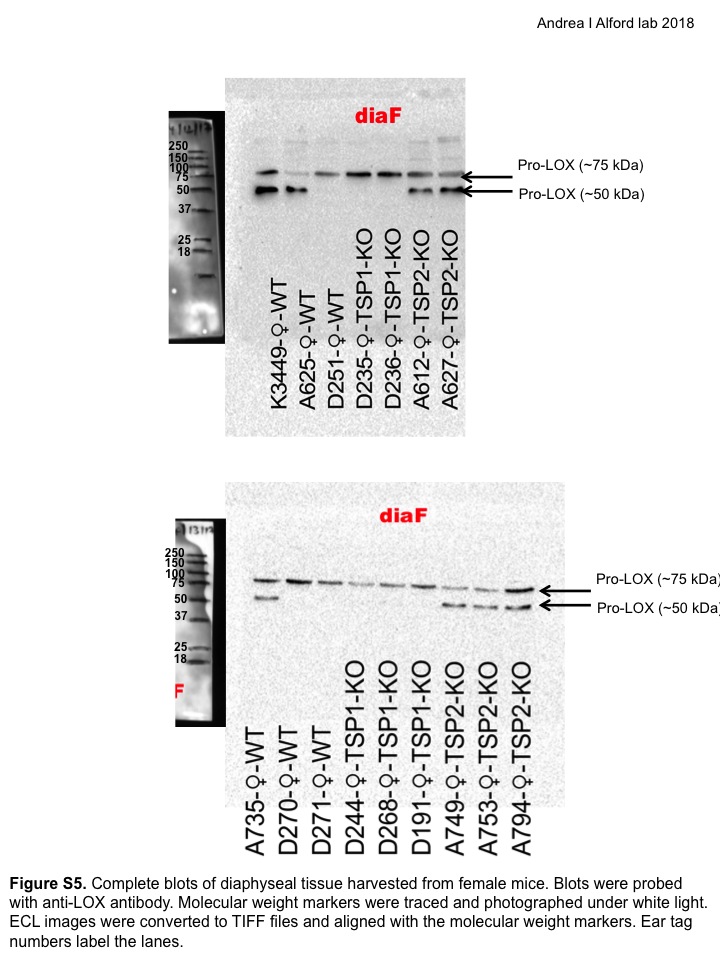

### Supplementary Materials

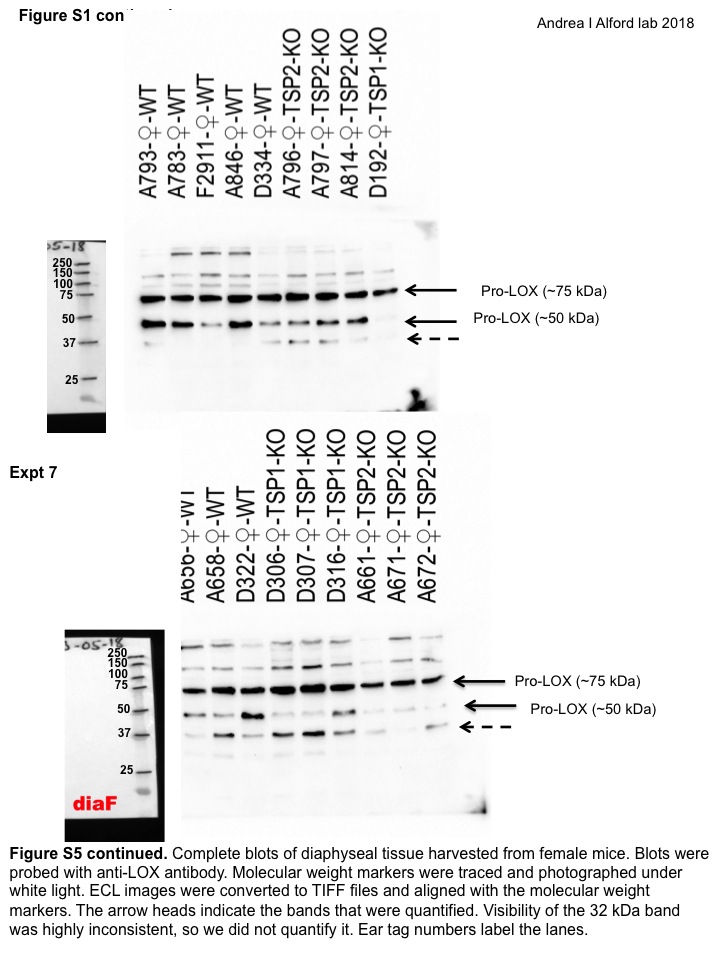

### Supplementary Materials

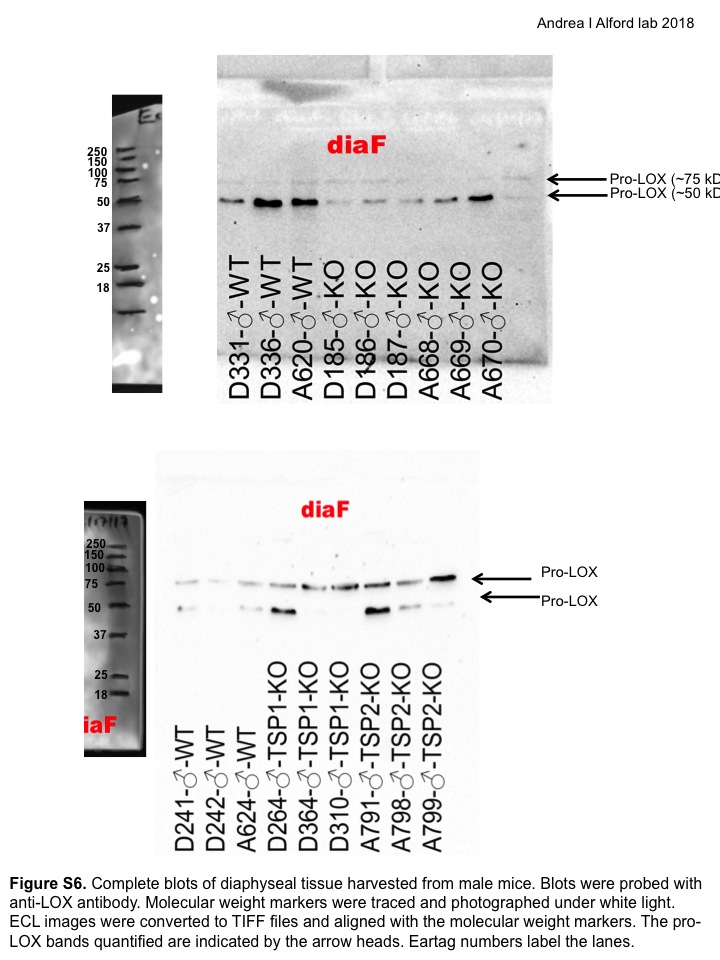

### Supplementary Materials

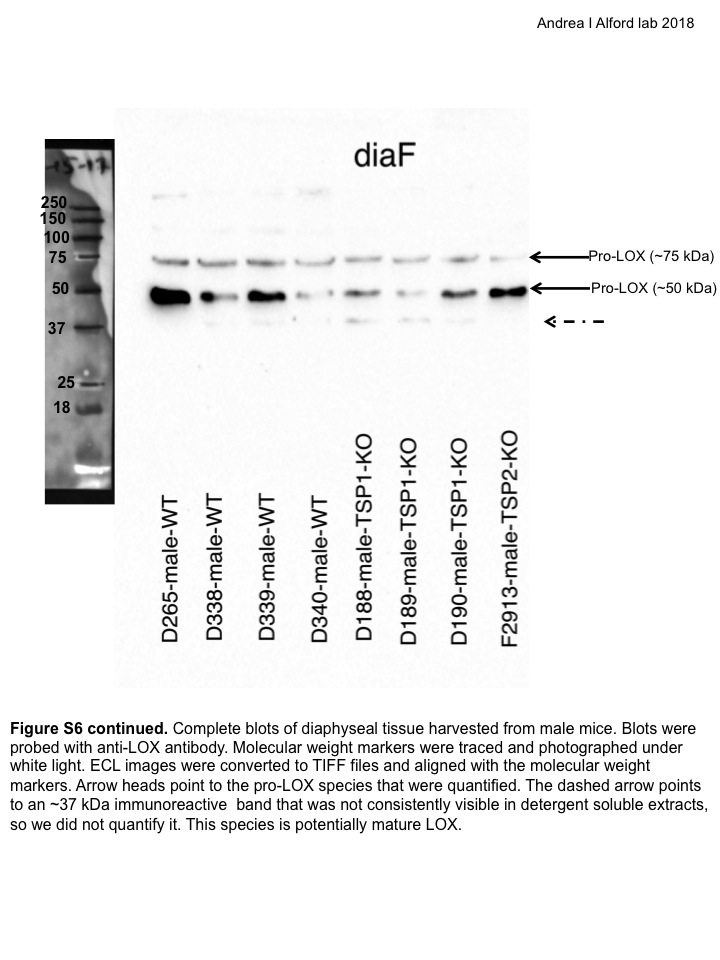

### Supplementary Materials

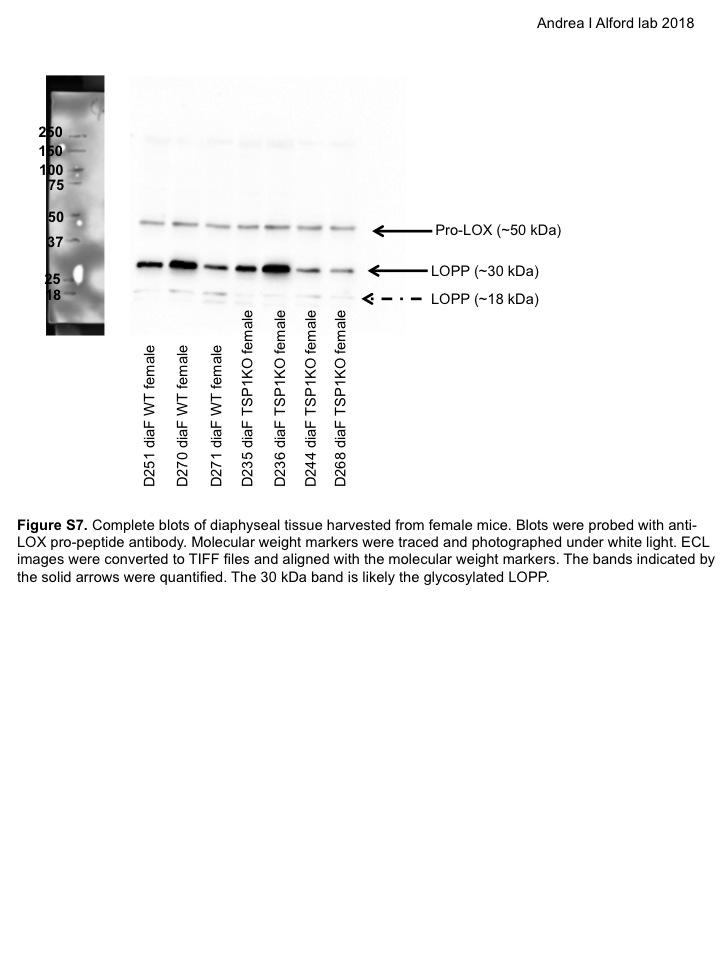
